# Spatial Chromatin Architecture Alteration by Structural Variations in Human Genomes at Population Scale

**DOI:** 10.1101/266981

**Authors:** Michal Sadowski, Agnieszka Kraft, Przemyslaw Szalaj, Michal Wlasnowolski, Zhonghui Tang, Yijun Ruan, Dariusz Plewczynski

## Abstract

This genome-wide study is focused on the impact of structural variants identified in individuals from 26 human populations onto three-dimensional structures of their genomes. We assess the tendency of structural variants to accumulate in spatially interacting genomic segments and design a high-resolution computational algorithm to model the 3D conformational changes resulted by structural variations. We show that differential gene transcription is closely linked to variation in chromatin interaction networks mediated by RNA polymerase II. We also demonstrate that CTCF-mediated interactions are well conserved across population, but enriched with disease-associated SNPs. Altogether, this study assesses the critical impact of structural variants on the higher order organization of chromatin folding and provides unique insight into the mechanisms regulating gene transcription at the population scale, among which the local arrangement of chromatin loops seems to be the leading one. It is the first insight into the variability of the human 3D genome at the population scale.

---

Around 20 million base pairs of a normal human genome (0.6%) are under structural variations, including deletions, duplications, insertions and inversions. This makes structural variants (SVs) the most prominent source of genetic variation among human individual genomes.

The potential malicious effect of SVs has been recognized but almost solely associated with altering gene copy number and gene structure – a number of studies relate copy number variants (CNVs) affecting gene regions to cancer (Malhotra and Sebat 2012), intellectual disabilities (Stankiewicz and Lupski 2010) and predispositions to various health problems (Zollino, et al. 2012), (Talkowski, et al. 2011). The vast majority of genetic variation occurs, however, in non-coding regions. Over 95% of single-nucleotide polymorphisms (SNPs) identified by genome-wide association studies (GWAS) are located outside coding sequences (Maurano, et al. 2012). Similarly, larger variants are significantly depleted in gene regions (Sudmant, et al. 2015).

Part of the SVs emerging in non-coding regions alter genomic loci recognized by proteins which organize the human genome in the cell nuclear space. Recent studies provided some insights into the impact SVs can have on spatial organization of the human genome. Examples of SVs altering borders of TADs in EPHA4 locus and causing pathogenic phenotypes by enabling spatial contacts between formerly isolated genomic functional elements were reported (Lupiáñez, et al. 2015). Positions of TAD boundaries were proven useful for inferring cancer-related gene overexpression resulting from variation in cis-regulatory elements (Weischenfeldt, et al. 2017). Accumulation of SVs proximal to TAD boundary occupied by CTCF was postulated to cause enhancer hijacking and PRDM6 overexpression in medulloblastoma samples (Northcott, et al. 2017). Hi-C maps were successfully used for detection of large-scale rearrangements, which were reported as frequent in cancer cells (Dixon, Xu, et al. 2018). Disruptions of chromosome neighborhoods were demonstrated – using CRISPR/Cas9 experiments – to activate proto-oncogenes (Hnisz, et al. 2016). An attempt was also made to model 3D chromatin structure including information on SVs and predicting enrichment/depletion of higher-order chromatin contacts caused by these variations (Bianco, et al. 2018). Efficacy of the modelling method in predicting SV-induced ectopic contacts at the level of TADs was shown for EPHA4 locus.

However, to our knowledge, there was no genome-wide systematic study on the impact of SVs on genome spatial organization analyzing the level of individual chromatin loops. One of the most recent reviews on the topic (Spielmann, Lupiáñez and Mundlos 2018) highlights the impact of SVs on genome spatial structure and the pathogenic potential of SVs altering the higher-order chromatin organization. Nonetheless, no attempt was made by the authors to assess what part of SVs emerging in normal human genomes cause functionally relevant chromatin spatial rearrangements and no genome-wide data is presented on how SVs influence the chromatin 3D architecture.

The recent advancements in chromosome conformation capture techniques, namely High-throughput Conformation Capture (Hi-C) (Lieberman-Aiden, et al. 2009), (Rao, et al. 2014) and Chromatin Interaction Analysis by Paired-End Tag Sequencing (ChIA-PET) (Fullwood and Ruan 2009), (Tang, et al. 2015) resulted in release of high-resolution chromatin interaction datasets. ChIA-PET, in particular, is able to capture individual chromatin contacts mediated by specific protein factors. In turn, the great effort of the 1000 Genomes consortium led to creation of the largest catalogue of human genomic sequence variations (Sudmant, et al. 2015) identified in over 2,500 human samples from 26 populations.

Taking advantage of the high quality ChIA-PET and population-scale SVs data, we discuss a mechanistic model of the impact of SVs on the chromatin looping structure, provide the first genome-wide analysis of this impact for the human genome and model SV-induced changes in 3D genomic structures observed in human population.

In our analyses of the impact of SVs on the 3D chromatin organization of the human genome, we pay a specific attention to chromatin interactions associated with enhancer regions and gene promoters. These interactions are likely to play a distinguished role in the regulation of gene transcription in a mechanistic fashion, bringing the genes and the regulatory elements close together in the nuclear space of the cell. We observe an interesting interplay between such genomic interactions and SVs.

## Results

### 3D genome

In this study we use ChIA-PET data as a representation of the higher-order spatial organization of the human genome. ChIA-PET generates high-resolution (∼100 bp) genome-wide chromatin contact maps. It identifies two types of chromatin interactions mediated by specific protein factors. The first type are highly reliable enriched interactions which appear in the data as closely mapped on the genome clustered inter-ligation paired-end-tag products (PET clusters). The second type are singletons, which reflect higher-order topological proximity (Tang, et al. 2015).

CTCF was shown to be the key protein factor shaping architecture of mammalian genomes (Ong and Corces 2014), (Rao, et al. 2014). We use ChIA-PET targeting on CTCF for GM12878 cell line, which is presently the most comprehensive ChIA-PET dataset for human. We inspected the anchoring sites of PET clusters from this dataset for the co-occupancy by CTCF and cohesin (RAD21 and SMC3 subunits), to select the set of high quality chromatin interactions mediated by CTCF in GM12878 cell (see Methods). We identified 42,177 such pairwise interactions. The median length of genomic segments joined by these interactions is 1,425 bp and 99% of them are shorter than 10 kb (Fig. 1f). Nucleotide sequences of these segments usually contain multiple CTCF motifs. Chromatin loops formed by the CTCF interactions have lengths in the order of 100 kb (Fig. 1f).

**Fig. 1.**
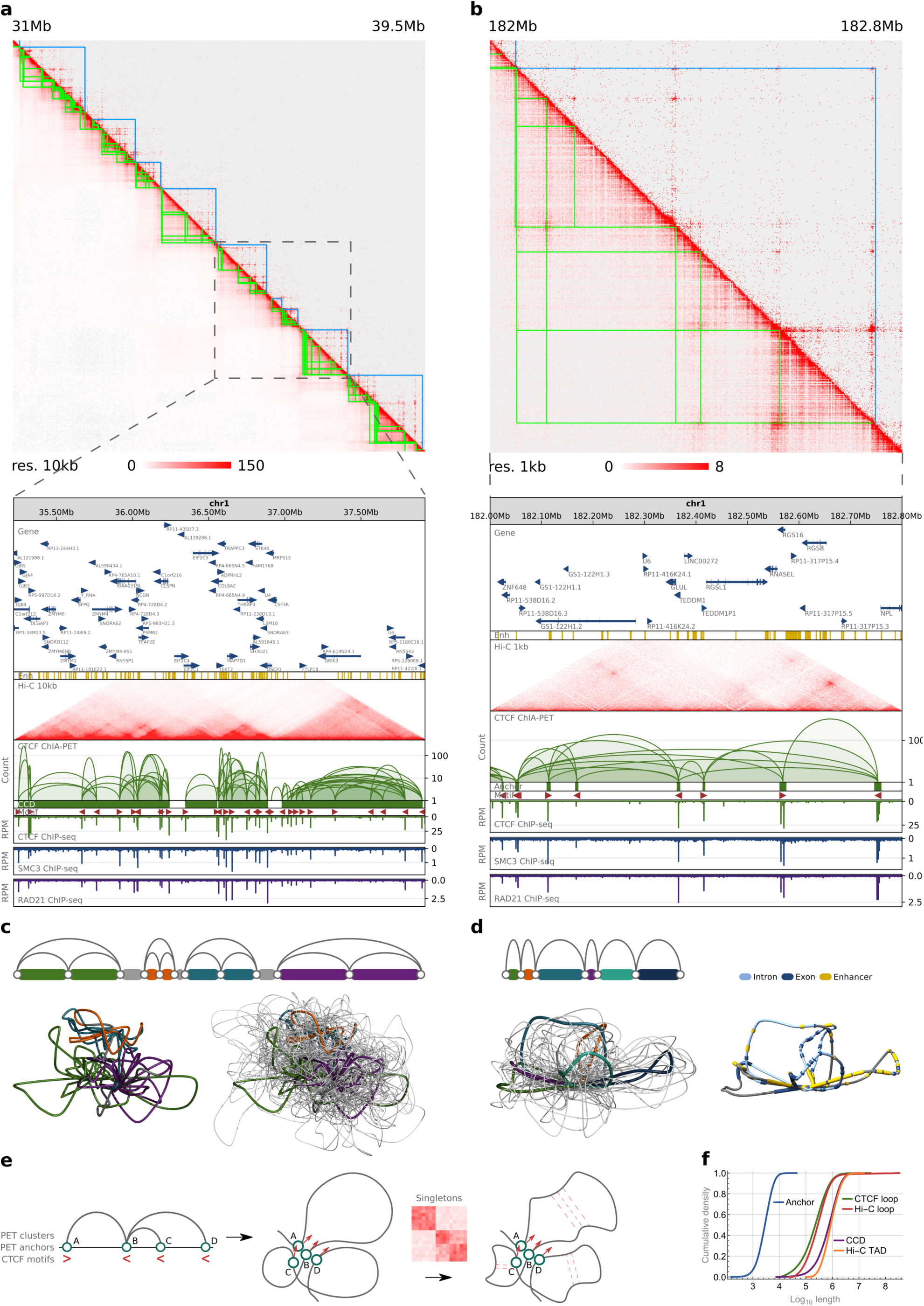
3D genome. Data aggregation and comparison. **a,** Comparison of Hi-C (lower-left) and ChIA-PET (upper-right) heatmaps of a 8.5 Mb genomic region in 10 kb resolution. Annotation for chromatin loops (green) and domains (blue) from ChIA-PET is presented on the heatmaps. The same ChIA-PET data is shown in a browser view together with ChIP-seq tracks for CTCF and cohesin subunits. The height of an arc indicates the strength of an interaction, which is measured by the number of clustered individual inter-ligation paired-end-tag products. Annotations of genes (GENCODE) and enhancers (ChromHMM) are presented. Arrows at genes mark direction of their transcription. **b,** Similar to (a), but a 0.8 Mb genomic region is presented in a heatmap in 1 kb resolution and in a browser view. **c,** 3D models of the genomic region marked in (a). For clarity, every topological domain in the region has a different color in the model. On the left, a single 3D model (centroid). On the right, an ensemble of 10 models with centroid model colored. Centroid is a structure most similar to all the others in an ensemble in terms of RMSD measure. **d,** 3D models of the genomic region shown in (b). Every loop has a different color. On the left, an ensemble of 10 structures. On the right, a single model (centroid) with functional genomic elements marked. **e,** Scheme presenting the chromatin modeling method at the level of loops. The method uses PET clusters, singletons and orientations of CTCF binding motifs to accurately model genome looping structures. **f,** Cumulative density plot showing the genomic span distribution of genomic structural elements identified in ChIA-PET and HI-C.

The interactions mediated by CTCF are not uniformly distributed over the genome but rather form highly interacting, predominantly hundreds of kilobases long chromatin blocks (chromatin contact domains, CCDs or topologically associating domains, TADs) separated by segments of weak and rare contacts (gaps). Based on the CTCF ChIA-PET data, genome of GM12878 cell was segmented into 2,267 CCDs (Tang, et al. 2015). The domains lengths vary from around 10 kb to few megabases with a median length of 750 kb. Only 1% of CCDs is longer than 2 Mb (Fig 1f).

Even though CTCF ChIA-PET captures only interactions mediated by the CTCF protein, it detects genomic structures and general structural features exhibited by the non-specific Hi-C data. On a global scale, whole-chromosome Hi-C and ChIA-PET contact maps are highly correlated (Spearman’s correlation coefficient in range 0.7 – 0.9) (Tang, et al. 2015). Locally, ChIA-PET and Hi-C heatmaps identify very similar landscape of genomic structures, both at the scale of topological domains (Fig. 1a) and chromatin loops (Fig. 1b). Borders of topological domains emerging from the CTCF ChIA-PET data are highly enriched in CTCF (Fig. 1a), similarly as it was shown for TADs identified in Hi-C (Rao, et al. 2014). We adopted coordinates of the outermost CTCF motifs identified in a CCD as indicators of its borders (see Methods). Length distributions of chromatin loops and topological domains called from ChIA-PET data are concordant with the respective statistics for Hi-C (Fig. 1f) (Tang, et al. 2015). All this indicates that CTCF ChIA-PET dataset generated for GM12878 cell is a high quality representation of the human 3D genome.

We investigated the directionality of CTCF motifs in anchors of the CTCF chromatin loops. 35,180 out of the 42,177 PET clusters had motifs of the unique orientation in both anchors. Among the 35,180 interactions we found 16,239 interactions with motifs in the anchors having convergent orientation (convergent loops), 12,195 interactions in tandem right orientation (tandem right loops), 4,090 tandem left loops and 2,656 divergent loops (see Methods).

We demonstrated that using CTCF ChIA-PET data a 3D model of an averaged genome structure which recovers architectural features of the genome can be built (Szałaj, et al. 2016). Chromatin models constructed with our computational tool (3D-GNOME) can be used to illustrate the most probable arrangement of genomic structural elements in 3D space: from chromosomes, through topological domains (Fig. 1c), to individual chromatin loops (Fig. 1d). The last two levels of the genome organization are in the main scope of this study, as topological domains are believed to be the structural units regulating gene transcription by spatially isolating groups of enhancers and genes (Lupiáñez, et al. 2015), (Rao, et al. 2014), (Tang, et al. 2015), (de Laat and Denis 2013). Spatial arrangement of these functional elements can also be visualized using our modeling method (Fig. 1d).

3D-GNOME is an optimization algorithm which returns models that fulfill spatial constraints coming from genomic interaction data. Typically, a number of solutions exists for a given set of constraints, and thus we present and analyze ensembles of 3D structures for every genomic segment modeled (Fig. 1c and 1d). 3D-GNOME uses PET clusters to position the binding sites relative to each other first, and then employs singletons, orientations of the CTCF motifs and biophysical constraints to model the shape of individual chromatin loops (Fig. 1e). The modeling algorithm includes all this information to make the models as accurate as possible at the finest chromatin scale. See Methods for description of the full modeling procedure at all levels of genome organization.

We complement the CTCF ChIA-PET dataset with the ChIA-PET targeting on RNA polymerase II (RNAPII) data generated for GM12878 cell in the same study (Tang, et al. 2015). It was postulated that multiple chromatin loops mediated by CTCF proteins form static macromolecular structures called interaction centers inside chromatin contact domains. Their anchors are enriched with active epigenomic markers. The interactions mediated by RNAPII are believed to draw cell-type specific genes toward interaction centers, thus having more functional and dynamic character than interactions mediated by CTCF (Tang, et al. 2015). Together, the ChIA-PET data of these two protein factors account for structural and functional aspects of the higher order organization and multiscale folding of chromatin 10 nm fiber in the human cell nucleus (Ou, et al. 2017).

### Genome topological variation

We demonstrated, by phasing CTCF PET clusters (see Methods), that allele-specific single-nucleotide variation in genome sequence can result in haplotype-specific chromatin topology (Tang, et al. 2015).

Differences in scores between CTCF binding motifs located on homologous chromosomes resulting from allele-specific SNPs (Fig. 2b and Supplementary Figure 1b) are reflected in relative values of CTCF signal. This can be observed for different lymphoblastoid cells. In case of GM12878, for which both datasets are available, differences in CTCF binding agree with presence/absence of chromatin contacts (Fig. 2a and Supplementary Figure 1a). The examples suggest that presence of particular chromatin loops in lymphoblastoid cells can be predicted based on their genotypes and genomic interactions identified in GM12878.

**Fig. 2.**
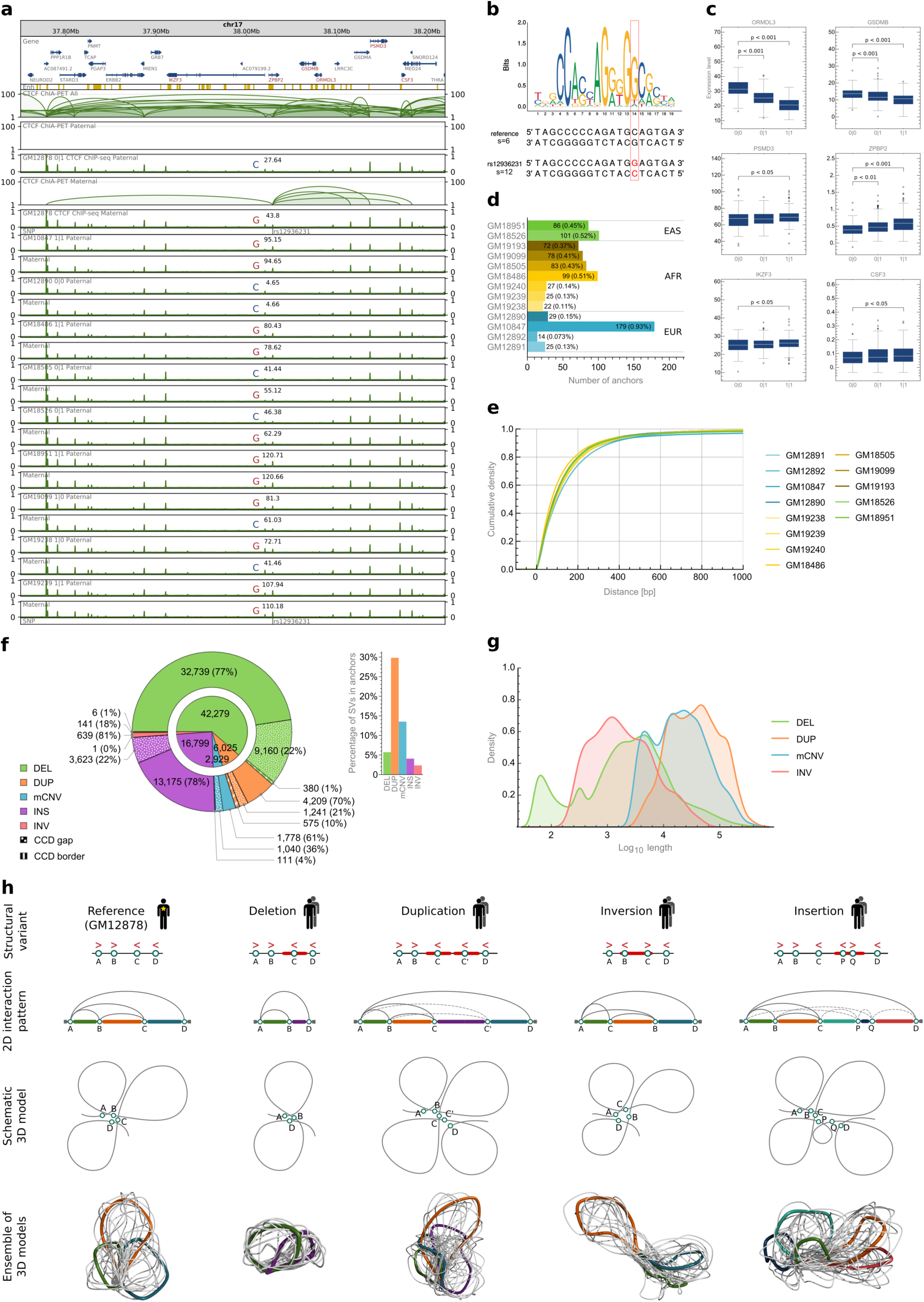
Impact of SVs on chromatin looping. **a,** Browser view of a 0.5 Mb genomic segment with asthma-associated SNP rs12936231 identified in a part of the human population. SNP rs12936231 alters sequence of a CTCF motif forming interactions. Haplotype-specific CTCF signals from 10 lymphoblastoid cells are presented along with haplotype-specific CTCF ChIA-PET interactions from GM12878. For each track, ChIP-seq signal values (originally in RPMs) were divided by the maximal value of the signal in the visualized region. Sum of the signal values over the genomic region occupied by the SNP-affected interaction anchor together with the genotype is marked in each signal track. **b,** Comparison of sequences and scores of CTCF binding motifs carrying the reference (C) and the alternative (G) alleles of rs12936231. **c,** Differences in gene transcription rates between genotypes set for rs12936231. Genes exhibiting differences in transcription which pass Mann-Whitney test with p-value < 0.05 were reported. **d,** CTCF anchors from GM12878 not intersected with CTCF ChIP-seq peaks identified in different lymphoblastoid cells. The anchors were filtered by consensus CTCF binding sites (see Methods). **e,** Cumulative density plot of distances between CTCF ChIP-seq peaks located in GM12878 interaction anchors and closest CTCF ChIP-seq peaks from different lymphoblastoid cells. **f,** Number of SVs, divided by type, intersecting (in case of interaction anchors), covering (in case of CCD boundaries) or contained in (in case of CCDs and CCD gaps) different genomic structural elements. **g,** Density plot showing genomic span distribution of SVs by type. **h,** Predicted impact of particular SV types on looping structure of the genome. Simplified chromatin looping patterns and 3D models are presented for the reference and its SV-altered versions.

Having shown how substitutions of single nucleotides can affect chromatin topology, here we concentrate on predicting how SVs impact genome looping organization. SVs are the major source of sequence variation among human genomes and given their larger size have a higher potential than SNPs to induce changes in chromatin folding. They were also shown to contribute more than SNPs to variation in gene expression among human samples (Chiang, et al. 2017).

We use ChIA-PET interactions collected from the GM12878 cell line as the reference network of chromatin spatial contacts in human lymphoblastoid cells.

Domain organization of chromatin is conserved across cell types and species (Dixon, Selvaraj, et al. 2012), (Dekker, et al. 2002), (Rao, et al. 2014). This indicates that chromatin contacts are also largely invariant across individuals. To assess the level of conservation of CTCF-dependent genome architecture across individuals we analyzed the abundance and arrangement of CTCF ChIP-seq peaks from 13 lymphoblastoid samples in genomic segments which were identified as CTCF interaction anchors in GM12878 cell line. The data originate from one study (Kasowski, et al. 2013) and include 5 samples of European ancestry (GM10847, GM12878, GM12890, GM12891, GM12892), 7 samples of African ancestry (GM18486, GM18505, GM19099, GM19193, GM19238, GM19239, GM19240) and 2 samples of East Asian ancestry (GM18526, GM18951). GM12878, GM12891, GM12892 and GM19238, GM19239, GM19240 are related. The datasets were rigorously filtered for comparisons (see Methods).

Our analysis shows that genomes from lymphoblastoid cells exhibit highly similar patterns of active CTCF binding sites. Over 99% of interacting anchors occupied by CTCF peaks in GM12878 cell is supported by CTCF peaks in each of the 13 other samples (Fig. 2d). Moreover, the overall distribution of CTCFs is shared among individuals – for about 90% of all CTCF peaks identified in anchors of PET clusters in GM12878 cell, a CTCF peak in each of the compared cells can be found within a distance of 400 bp (Fig. 2e). CTCF interacting anchors and borders of CCDs identified in GM12878 cell are highly enriched with CTCF ChIP-seq peaks found in the other 13 lymphoblastoid cells (Supplementary Figure 2).

Since every chromatin segment carrying a CTCF has a potential to create an interaction with another segment having a CTCF attached to it, we claim that highly similar patterns of CTCF distribution along lymphoblastoid genomes indicate that they exhibit highly similar patterns of chromatin interactions mediated by CTCF.

Similar analyses were performed for RNAPII interacting anchors using RNAPII ChIP-seq peaks (Supplementary Figure 3). Significantly bigger but moderate differences among samples were observed for this data.

Having tested the overall resemblance of 3D genomes in lymphoblastoid cells, we match SVs detected in human population by the 1000 Genomes Project (Sudmant, et al. 2015) with the reference network of chromatin interactions to obtain individualized chromatin interaction patterns and assess topological variability among human genomes.

There are 68,818 unique structural variants deposited in the 1000 Genomes Catalogue of Human Genetic Variation (CHGV), including deletions, duplications, multiallelic CNVs, inversions and insertions (Fig. 2f). 44% of them are shorter than 1 kb and only 22% is longer than 10 kb (Fig. 2g).

Most of the SVs reside inside CCDs, not intersecting borders of domains nor CTCF interacting anchors (Fig. 2f). However, the rate of duplications overlapping genomic structural elements is distinctively high.

While the SVs that miss the interacting binding sites in most cases have limited impact on the final structure (resulting only in shortening, or extending of the corresponding chromatin loops), the SVs that overlap the interacting sites may partially modify the interaction pattern and in turn cause serious changes of the 3D structure (Fig. 2h). Specifically, deletion removes an interacting anchor therefore deleting all chromatin loops mediated by this genomic site; duplication introduces new interaction site, which has the same underlying sequence specificity to form chromatin loops as the original duplicated site; inversion which encircles a CTCF binding motif will revert its directionality therefore affecting the chromatin looping of the neighboring region; and finally, insertion containing CTCF motif enables new interaction sites capable of forming chromatin loops with other CTCF binding sites.

We extended our modeling approach, 3D-GNOME, to include information on SVs in recovery of 3D chromatin structures (see Methods). Our algorithm models individual chromatin loops, meaning that the remodeling effect of a genetic variant disrupting a single pair of interacting genomic segments will be represented in the model (Fig. 2h).

### Topological impact of structural variations

CTCF ChIP-seq data confirms that there are SVs which result in altered activity of reference interaction anchors. As an example, deletion chr14:35605439-35615196 of an interaction anchor leads to a significant depletion of CTCF signal in heterozygous samples and even to a complete vanishing of the signal in a homozygous sample (Fig. 3a). The CTCF signal drop reflects the lower or no potential of CTCF to bind to this segment. Therefore, in a cell line exhibiting the deletion, all of the chromatin contacts formed by this locus would not be present in one or both of the homologous chromosomes, depending on the genotype (Fig. 3b). The deletion is located in an intron of gene KIAA0391, but does not excise any coding sequence. Nevertheless, the genotypes show statistically significant differences in transcription rates of several genes (Fig. 3d). Deletion chr14:35605439-35615196 removes chromatin contact bringing NFKBIA and one of the enhancers together in 3D space, leading to a different arrangement of enhancers around the gene (Fig. 3c). Even though the landscape of functional elements around NFKBIA gene is complex to the extend, that refrains from drawing definite conclusions, the deletion significantly reduces access of the gene promoter to enhancers (Fig. 3e), which explains the depletion of NFKBIA expression observed for samples carrying it. The deletion causes complex spatial rearrangements also around other genes, which contribute probably to the differences in their transcription rates between samples of different genotypes.

**Fig. 3.**
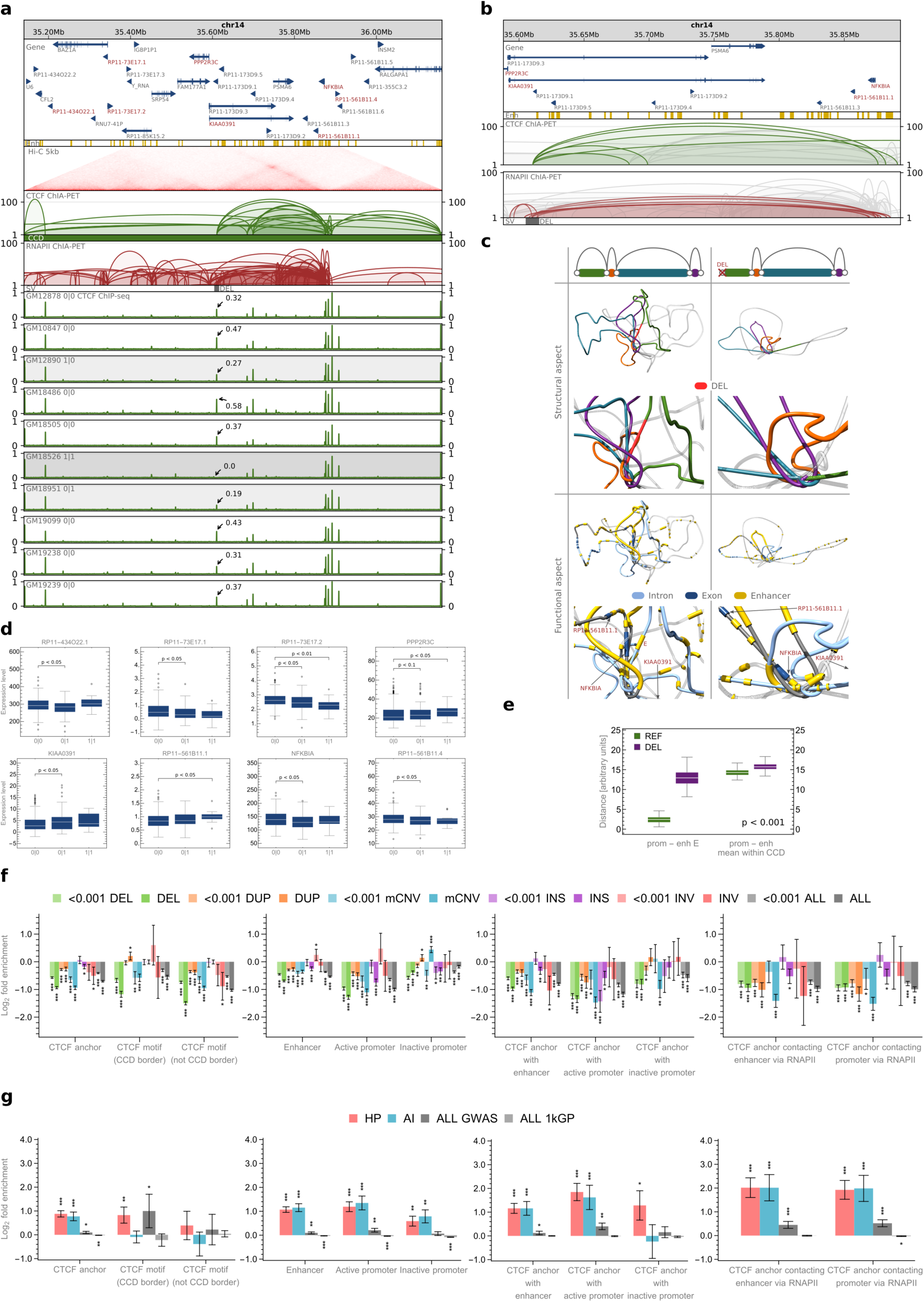
Impact of SVs on genome organization at the population scale. |**a,** Browser view of a 1 Mb genomic segment with a deletion identified in a part of the human population. The deletion removes a CTCF anchor with enhancer (E) located in an intron of KIAA0391. CTCF ChIP-seq signals from 10 lymphoblastoid cells of different genotypes are presented for comparison. For each track, ChIP-seq signal values (originally in RPMs) were divided by the maximal value of the signal in the visualized region. The highest signal peak in the genomic region covered by the deletion is marked in each signal track. **b,** Close-up on the chromatin interactions affected by the deletion. **c,** 3D models of the domain shown in (a) with and without the deletion. Loops shown in (b) are colored, as presented on schematic drawings. The spot of the variation is zoomed in separate pictures. Every picture has its replicate, in which genes and enhancers are marked. Selected genes are signed (arrows are pointing towards the TSSs). **d,** Differences in gene transcription rates between genotypes set for the deletion. Genes exhibiting differences in transcription which pass Mann-Whitney test with p-value < 0.05 were reported. **e,** Euclidean distance between the NFKBIA promoter and enhancer E and mean distance between the promoter and enhancers located in the same CCD. In green, distribution of distances calculated in 3D models of the reference structure (REF), in magenta – in models with the deletion introduced (DEL). For each case 100 models were generated. The differences between REF and DEL groups are significant (p-values < 0.001). **f,** Enrichment/depletion of genomic structural elements with SVs of different types and of different VAF (VAF < 0.001 and VAF ≥ 0.001). In case of CCD borders, only these fully imbedded in SV intervals are counted as affected, whereas for other structural elements ≥1 bp overlaps are counted. **g,** Enrichment/depletion of genomic structural elements with the 1000 Genomes Project SNPs, all GWAS SNPs, GWAS SNPs associated with hematological parameters (HP), and with autoimmune diseases (AI).

On the other hand, duplications of CTCF-mediated interacting genomic segments result in distinctively high relative values of CTCF signal in those segments in affected samples (Supplementary Figure 4a and 4b). The signal enrichment caused by those duplications supports a hypothesis that they create additional CTCF-binding loci with the potential to form additional long-range genomic contacts in the affected genomes.

Inversions in CTCF binding sites also modulate CTCF signal (Supplementary Figure 5a), which indicates they introduce changes in chromatin looping.

Genotypes set by such SVs exhibit significant differences in expression of particular genes (Fig. 3d, Supplementary Figures 4c and 5b).

Apart from SVs disrupting long-range chromatin interactions that join genomic segments located within one topological domain, there are examples of SVs modifying domain boundaries. Deletion chr1:109366972-109372000 is one of them (Supplementary Figure 6a). Samples genotyped by this deletion demonstrate statistically significant differences in transcription rates of PRPF38B gene (Supplementary Figure 6b).

An aggregate analysis (see Methods) shows that CTCF ChIP-seq signals from samples having deletions (duplications) which intersect CTCF interacting anchors are depleted (enriched) in those anchors compared to signals from samples with the reference genotype (Supplementary Figure 7).

To statistically assess the impact of SVs on the spatial organization of the genome, we analyzed their positions in relation to genomic structural elements like anchors of PET clusters, borders of topological domains, or gaps between the domains.

We observe that anchors of CTCF PET clusters are depleted of SVs (Fig. 3f) and that the rate of depletion is consistent among loops of different directionality (Supplementary Figure 8). Interestingly, CTCF interacting anchors that are not supported by cohesin peaks are slightly enriched with rare inversions, which is not the case for anchors occupied by cohesin (Supplementary Figure 9). This may suggest that genomic sites forming CTCF-mediated interactions, which are not stabilized by cohesin, are specifically prone to motif reorientation.

We further identified anchors of CTCF PET clusters intersected with enhancers and active and inactive gene promoters (see Methods). These anchors have a distinguished potential to play an important role in gene regulation. It appears as a general rule (applicable to functional regions located both inside and outside CTCF anchors) that genomic variations target enhancers and inactive promoters more frequently than active promoters (Fig. 3f). More interestingly, we observe that enhancers and promoters located in anchors of CTCF PET clusters are significantly more conserved than the respective functional regions residing outside the CTCF anchors (Fig. 3f). This indicates the importance of the genomic architecture mediated by CTCF in proper genome regulation. We additionally examined the conservation of CTCF anchoring sites, which are in contact with enhancers and gene promoters via RNAPII ChIA-PET interactions and they also seem to be more conserved than the respective genomic functional elements (Fig. 3f).

Surprisingly, borders of CCDs do not seem to be distinctively well conserved. CTCF binding motifs we identified in CTCF ChIP-seq peaks outside CCD borders are significantly more depleted of SVs than the CTCF motifs indicating borders of CCDs (Fig. 3f). Moreover, CCD borders are targets of many duplications (Fig. 2f and 3f). There is also a slight enrichment of rare inversions in CCD borders, but because the set of inversions is small the result is not statistically significant. However, we hypothesize that inverting the CTCF motifs at the borders of topological domains can be an important mechanism of genome reorganization and regulation. 6 out of 786 inversions from the CHGV switch the directionality of CCD borders (Fig. 2f). Inversion chr20:39359355-39379392 is one of them (Supplementary Figure 10a). Interestingly, it was found only in genomes of Africans and Puerto Ricans and correlates with transcription rates of a pair of neighboring genes (Supplementary Figure 10b).

Stronger conservation of CTCF anchors intersected with and connected to the known enhancers and promoters as compared to the conservation of enhancers and promoters located outside CTCF anchors suggests that mutations of these anchors may lead to serious deregulations of gene expression and can be related to disease. To test this hypothesis, we intersected CTCF anchors with SNPs previously associated with disease in GWAS. Having in mind the type of cell we examine, we created separate sets of GWAS SNPs associated with hematological parameters and autoimmune diseases. Our analysis indeed shows a significant enrichment of these SNP classes in CTCF anchors intersected with enhancers and active promoters (Fig. 3g). Particularly, the enrichment is high in CTCF anchors being in RNAPII-mediated contact with enhancers and promoters. Both former and latter type of anchors are enriched with all GWAS SNPs. Importantly, enhancers and active promoters, which are not intersected with CTCF anchors, are notably less enriched with GWAS SNPs than CTCF anchors associated with these functional elements (Fig. 3g). Generally, CTCF anchors and CCD boundaries are enriched with GWAS SNPs (Fig. 3g). Our observations are consistent with the studies on capture Hi-C (Mifsud, et al. 2015), (Martin, et al. 2015), and are additionally highlighting the role of CTCF in shaping the network of functionally important genomic contacts.

We investigated particular examples of SNPs associated with autoimmune diseases (rheumatoid arthritis and vitiligo, rs4409785 T/C) and hematological parameters (red blood cell distribution width, rs57565032 G/T) (Supplementary Figures 11a and 12a). Both alter strongest CTCF binding motifs in the corresponding interaction anchors. However, their effect on CTCF binding is opposite: rs4409785 increases strength of CTCF motif it modifies (Supplementary Figure 11b), rs57565032 decreases (Supplementary Figure 12b). It is reflected in CTCF signals corresponding to different genotypes (Supplementary Figures 11a and 12a). No other SNPs affect CTCF motifs in those interaction anchors in presented genomes. Samples genotyped by these SNPs demonstrate significant differences in transcription rates of particular genes (Supplementary Figures 11c and 12c). One of them, MAML2, has been associated with cancer traits.

The already presented SNP rs12936231 (Fig. 2a) has been reported as a high-risk allele for asthma and autoimmune diseases and suggested to cause chromatin remodeling and alter transcription of certain genes, including ZPBP2, GSDMB and ORMDL3 (Verlaan, et al. 2009). We also found correlation of genotypes set by rs12936231 with transcription rates of ZPBP2, GSDMB and ORMDL3 (Fig. 2c). Gene IKZF3, which also exhibits a correlation with rs12936231, has been related to B-cell chronic lymphocytic leukemia.

Examples of SNPs in interacting anchors, but not associated with disease so far, can also be found. SNP rs60205880 alters CTCF-mediated chromatin looping and transcription of certain genes (Supplementary Figure 1). One of them, CCDC19, has been associated with bilirubin levels; another, IGSF8, is a member of an immunoglobulin superfamily. This demonstrates the potential of investigating SVs that affect genomic structural elements for identification of health-affecting genetic variation and mechanisms by which SVs are related to disease.

The important question is how large the structural variation among healthy individuals is. Individuals from the 1000 Genome Project have from 2,571 to 6,301 SVs, which affect from 1,024 to 1,419 CCDs and 52 – 330 CTCF anchors (Supplementary Figure 13). Almost all CCDs (98%) have an overlap with at least one SV from the CHGV. However, serious changes in local genome architecture are introduced by disruptions of interaction anchors rather than modifications of genomic regions between them. We identified 4,845 unique patterns of SVs altering interaction anchors in CCDs (we treat 2 patterns as identical if anchor-intersecting SVs they contain are the same; patterns are limited to single CCDs). Together with the 2,267 reference CCDs it gives the number of CTCF-mediated topologies of genomic domains occurring in the 1000 Genomes Project population. We note that types of SVs are well separated in those patterns (Supplementary Figure 14). 87% of the patterns are comprised of only one SV type. There are 1,500 patterns consisting of 2 or more SVs and in 878 (59%) of them all SVs are of the same type.

### Population-specific topological alterations affected by structural variations

We additionally analyzed intersections of SVs with genomic structural elements in the context of five continental groups: Africa (AFR), the Americas (AMR), East Asia (EAS), Europe (EUR) and South Asia (SAS). These populations are defined by the 1000 Genomes Project (The 1000 Genomes Project Consortium 2015) (Supplementary Table 1).

In all the populations, deletions of interacting anchors are more frequent than duplications (Supplementary Figure 16a). This is not true for CCD borders (Supplementary Figure 16b), which agrees with the previous analyses showing that CCD borders are enriched with duplications (we note that there are significantly more deletions than duplications in the set of detected SVs (Fig. 2f)). Alterations of topological domain boundaries can be a general mechanism of genome structure evolution. The above results suggest that such generic mechanism – similarly to the evolutionary process of introducing gene alterations by duplications – could use redundancy as a security measure. It could leave one chromatin loop with the original transcriptional function under evolutionary pressure, whereas second could be acquiring novel local spatial landscape for genes and regulatory elements.

Our analysis shows that individual genomes from populations of African ancestry have the largest number of deletions in CTCF interaction sites (Supplementary Figure 16a). This is partly due to an outstanding number of all deletions identified in those genomes (Supplementary Figure 15). However, we still observe that African genomes, together with European genomes, have CTCF-anchor sites less depleted of SVs unique to populations than genomes of other ancestries (Supplementary Figure 16c).

Interestingly, SVs found only in European genomes are significantly less depleted in CTCF interaction anchors intersected with enhancers or gene promoters than SVs unique to the rest of 5 distinguished continental groups (Supplementary Figure 17). We observe the same effect for borders of CCDs (Supplementary Figure 16c), but not for enhancers and promoters residing outside CTCF anchors (Supplementary Figure 18). This may suggest that part of the SVs identified in non-European populations overlap interaction sites specific for these populations and not observed in the reference 3D genome.

South Asian genomes, on the other hand, have distinctively large number of duplicated CTCF-anchor (Supplementary Figure 16a) and CCD-border (Supplementary Figure 16b) sites. Whereas the distinctive number of altered structural elements in genomes of African ancestry could be expected based on the large genomic sequence variability in this population reported earlier (The 1000 Genomes Project Consortium 2015), high structural variability in populations of South Asia is surprising. The ethnic groups which raise the statistics for South Asian continental group, especially those related to CNVs, are: Indian Telugu in the UK (ITU), Punjabi in Lahore, Pakistan (PJL) and Sri Lankan Tamil in the UK (STU) (Supplementary Figures 16f and 16g). As a comparison, corresponding statistics for African and European continental groups seem to be more stable across ethnic groups (Supplementary Figure 19). To investigate this further, we analyzed homozygous SVs. We hypothesized that the elevated number of structural changes observed in South Asian genomes could be caused by the high number of homozygous SVs that some of the populations in this continental group exhibit due to high consanguinity rates (Bittles, et al. 1991).

There are 13,767 homozygous SVs in the CHGV (we treat a CNV as homozygous, when there is a non-reference copy number on both homologous chromosomes). According to the data, genomes from East Asia, not South Asia, carry the largest number of the homologous SVs (Supplementary Figure 16h). However, the differences in homozygous sequence variation are not reflected in the number of homozygously altered CTCF anchors. The latter seems not to be changing across populations (Supplementary Figure 16i).

The fruitful study of natural human knockouts performed on a cohort of 10,503 Pakistanis by the Human Knockout Project (Saleheen, et al. 2017), made us investigate the homozygous SVs from the CHGV identified uniquely in a single population. We took the 1,317 knocked out genes found in individuals from South Asia (in majority belonging to Urdu and Punjabi ethnic groups, over 70%) (Saleheen, et al. 2017) and considered 656 CCDs they were located in. It turns out that homozygous SVs identified RNAPII anchors in the largest number of CCDs (10 and 11 respectively) containing the gene knockouts (Supplementary Figure 16k), even though a moderate number of population-specific homozygous SVs was found for this group (Supplementary Figure 16j). This suggests that gene knockouts may be accompanied (preceded, followed or assisted) by homozygous structural rearrangements.

For each of the continental groups we prepared a list of patterns (similar to these described in the previous section) of anchor-intersecting SVs, which alter CCDs in the individuals belonging to this group. Even though most of the patterns are population-specific, we found 305 (6%) patterns common for all 5 continental groups (Supplementary Figure 16e). CCDs in which we found the common SV-patterns are characterized by particularly high number of gene promoters, including promoters of housekeeping genes (Supplementary Figure 16d). There are statistically more gene promoters in those CCDs than in other CCDs with modified anchors and in domains covering segments without changes in CTCF-anchor sites. It is worth noticing that more promoters are located in CCDs containing CTCF anchors under variation than in those without them, which may suggest that architecture of transcriptionally active genomic regions is more prone to mutation. CCDs with CTCF anchors under rare variation contain statistically more promoters of active genes than CCDs with CTCF anchors affected only by frequent SVs (Supplementary Figure 16d). Moreover, rare SVs happen to affect CTCF anchors in domains containing outstanding number of promoters (Supplementary Figure 16d). CCDs with rare variants in CTCF interacting anchors can have up to 96 promoters of active genes (compared to 38 in CCDs with frequent SVs in anchors).

### Regulation of gene transcription altered by topological variations in population

By combining information on chromatin interactions and population-scale genetic variation with transcriptome data from 462 lymphoblastoid cell lines gathered by gEUVADIS consortium (Lappalainen, et al. 2013), we were able to draw first to our knowledge population-scale evidence-supported conclusions on the functional relation between SVs and genome architecture and provide a deeper insight into the functional role of genetic variation in the human genome. The results of our analysis indicate that SVs influence gene transcription primarily by rearranging local looping structure of the genome.

For 445 out of the 462 samples provided by gEUVADIS consortium there are also genotypes available in the 1000 Genomes Project database. We thus used PEER normalized (Lappalainen, et al. 2013) gene expression levels of these 445 individuals for association with their genotypes to identify expression quantitative trait loci (eQTLs). We use the term eQTL for any variation of genomic sequence which is identified as having effect on the gene transcription level. The eQTL analysis was performed only with SVs, excluding SNPs.

We performed Principal Component Analysis (PCA) on the expression data, which pinpointed 14,853 genes having the biggest variation in expression rates among individuals from all 23,722 genes present in the gEUVADIS dataset. We then related expression levels of each of the 14,853 genes to genotypes (see Methods).

In the studies on eQTLs published so far (Sudmant, et al. 2015), (Lappalainen, et al. 2013), (Schlattl, et al. 2011), (Stranger, et al. 2007), (Pickrell, et al. 2010), (Veyrieras, et al. 2008) a genomic region of arbitrary size around a gene in question was conventionally set, and only genetic variants located within this linear region were tested for being eQTLs for this gene. We argue that a more natural approach is to look for eQTLs within the whole topological domain the gene is located in. Therefore, for each of the selected genes, we evaluated associations of its expression levels with all the genotyped SVs residing in the same CCD. For every gene-SV pair, least-square linear regression was performed and the significance of the slope was then tested in the permutation test. The resulting *P-*values were adjusted for multiple testing to control the False-Discovery Rate (FDR). We set a threshold of acceptance for FDR ≤ 10% (see Methods).

We identified 234 unique SV-eQTLs modifying expression levels of 192 genes. The majority of the eQTLs found (55%) are deletions (Fig. 4a).

**Fig. 4.**
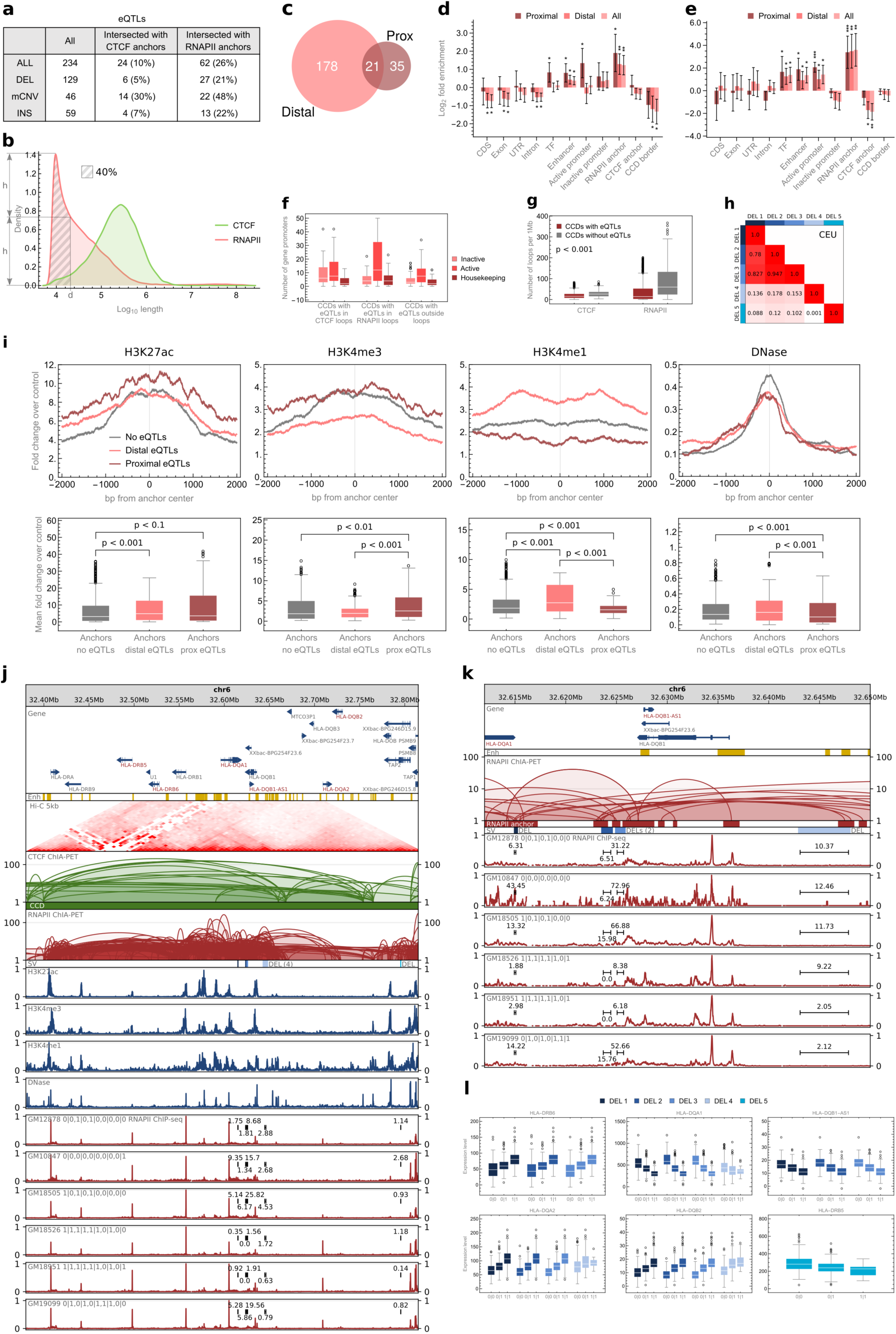
Role of chromatin rearrangements in regulation of gene transcription. **a,** Table summarizing identified eQTLs and their intersections with interaction anchors. **b,** Density plot showing genomic span distribution of PET clusters. *d* is the value (17,800 bp) by which eQTLs were split into proximal and distal. **c,** Venn diagram showing number of proximal (Prox) and distal eQTLs. **d,** Enrichment/depletion of genomic elements with eQTLs. **e,** Enrichment/depletion of genomic elements with eQTLs of housekeeping genes. **f,** Abundance of gene promoters in CCDs, in which eQTLs were identified. **g,** Distributions of chromatin loop density in CCDs in which eQTLs were identified and in other CCDs. The density is measured for a particular CCD as an average number of CTCF-/RNAPII-mediated chromatin loops covering a 1 Mb fragment of this CCD. Differences between the groups are significant (p-values < 0.001). **h,** Linkage disequilibrium (measured as r^2^ value in the CEU population) between deletions shown in (j) and (k). Colors are assigned to the deletions as in (j), (k) and (l). **i,** Signal strength of histone marks and DNase hypersensitivity sites in interaction anchors intersected with proximal eQTLs, distal eQTLs and not intersected by eQTLs. For each mark two plots are presented. A signal track around anchor center (±2 kb) showing values for each genomic position averaged over all anchors from a given group. A boxplot showing mean signal values in the same regions. Original signal values represent fold change over control. CTCF and RNAPII anchors were analyzed jointly. **j,** Browser view of a 0.4 Mb genomic segment with 5 deletions identified in a part of the human population, which disrupt RNAPII anchors and are eQTLs for 6 neighboring genes (signed with the red font). Each deletion has its color. RNAPII ChIP-seq signals from 6 lymphoblastoid cells of different genotypes are presented for comparison. For each track, normalized ChIP-seq signal values were divided by the maximal value of the signal in the visualized region. Sum of the signal values over the genomic regions occupied by the deletions is marked in each signal track. H3K27ac, H3K4me3, H3K4me1 and DNase-seq signal tracks from GM12878 are shown. **k,** Close-up on the RNAPII-mediated interactions affected by 4 of the 5 deletions. Only loops affected by the deletions are shown for clarity. **l,** Genes which transcription is correlated with one or more of the deletions shown in (j) and (k) (p-value < 0.001). Boxes with transcription rates associated with a particular deletion are marked with the color assigned to the deletion, as in (j) and (k).

Earlier studies on eQTLs were limited in exploring the causal relation between genetic variation and gene expression to analyzing gene-variant and exon-variant intersections (Schlattl, et al. 2011), (Pickrell, et al. 2010), or the influence of genetic variation on transcription factor binding sites (TFBSs), transcription start sites (TSSs) or transcription end sites (TESs) (Lappalainen, et al. 2013), (Veyrieras, et al. 2008), (Gaffney, et al. 2012). In particular, one of the latest to our knowledge big study on the impact of structural variation on human gene expression, reported that over 88% of predicted causal SVs did not alter gene structure or dosage (Chiang, et al. 2017). The study showed enrichment of causal noncoding SVs in regions occupied by transcription factors or surrounding genes at distances up to 10 kb, but no deepened analysis of these regions was performed. Our analysis gives a broader idea of this relation and sheds light on the mechanisms through which SVs take part in genome regulation.

In agreement with (Veyrieras, et al. 2008), (Gaffney, et al. 2012), (Chiang, et al. 2017) we observe enrichment of eQTLs in TFBSs but we see significantly higher enrichment of these in genomic regions responsible for chromatin spatial organization. We divided the identified eQTLs into two sets: those located on the DNA chain closer than 17,800 bp to the genes they modify (proximal) and those located further apart (distal) (see Methods). The splitting value of 17,800 bp was chosen based on the distribution of RNAPII PET clusters lengths. It is a value for which the density of lengths of RNAPII clusters is equal to the half of the maximum density (Fig. 4b). The division is not exclusive – some of the eQTLs correlated with more than one gene are distal for one of them and proximal for other (Fig. 4c).

Active promoters and TFBSs are enriched with proximal eQTLs demonstrating their importance as gene-adjacent regulatory sites. Enhancers apart for being enriched with proximal eQTLs are enriched with the distal ones and represent regulatory elements interacting with genes by the nuclear space. However, the genomic elements most enriched with both proximal and distal eQTLs are anchors of RNAPII PET clusters (Fig. 4d). The abundance of eQTLs in the anchoring regions of the strong chromatin interactions mediated by RNAPII reaffirms the crucial role of this element of genome architecture in gene regulation.

As an example of eQTLs altering chromatin looping mediated by RNAPII, we investigate 5 deletions located in a HLA region (Fig. 4j). All of these deletions affect RNAPII interacting anchors (Fig. 4k) and are correlated with one or more of 5 HLA genes neighboring them (Fig. 4l). Three of the deletions are in a very strong linkage disequilibrium (LD) with each other (tested on the Central European population, Fig. 4h).

The concept of CTCF foci given in (Tang, et al. 2015) suggests that CTCF interactions organize genome into structural interaction centers, within which RNAPII acts as a coordinator drawing genes into contact with center’s focal point where transcription happens. We hypothesize that proximal eQTLs modify TFBSs, TSSs and TESs of genes as well as gene sequences but mostly they alter genes’ spatial contacts with regulatory elements and possibly with transcription foci, which has an immediate and straightforward influence on genes’ expression levels. Distal eQTLs have in turn higher potential to, apart from altering long-range RNAPII interactions, disrupt CTCF interactions that are longer than RNAPII-mediated chromatin loops (Fig. 4b) and shape the spatial structures of the whole topological domains.

Deletion chr1:248849861-248850138 is an example of eQTL disrupting chromatin looping mediated by CTCF. It removes the strongest CTCF binding motif in an interacting anchor containing an enhancer (Supplementary Figure 20a) and is eQTL for 5 genes residing on a chromatin loop formed by the anchor (Supplementary Figure 20b). Three of them (OR2T3, OR2T10 and OR2T34) are members of the olfactory receptor gene family.

Another example of identified eQTL which alters CTCF-mediated chromatin structure, is duplication chr17:44341412-44366497 (Supplementary Figure 21a). It duplicates border of a CCD and its emergence correlates with transcription of a pair of genes (Supplementary Figure 21b).

Even though we observed examples, we did not find anchors of CTCF PET clusters to be enriched with distal eQTLs (Fig. 4d). The fact we note, however, is that 16 of the identified eQTLs (23% of the anchor-intersecting eQTLs) intersect both RNAPII and CTCF anchors (Fig. 4a) and 35 (56%) of the eQTLs intersecting RNAPII anchors were detected in CCDs in which eQTLs targeting CTCF anchors were also found. This suggests that a change in gene expression observed among individuals can often be a result of a coordinated modification of RNAPII and CTCF anchors, but more investigation is needed to confirm this claim. Interestingly, CCDs in which eQTLs alter RNAPII anchors tend to embrace more active genes and housekeeping genes than CCDs with eQTLs not overlapping any interacting segments (Fig. 4f). On the other hand, CCDs with eQTLs in CTCF anchors contain many inactive genes (Fig. 4f).

Furthermore, we suspect that the enrichment analysis does not indicate that alterations of CTCF anchors significantly contribute to variation of gene expression in population because the disruption of CTCF chromatin contacts would often provoke drastic changes in local spatial organization of a genome not observed in healthy people. As we showed earlier, SNPs associated with disease favorably emerge in CTCF anchors (Fig. 3g).

For comparison with Capture Hi-C (CHi-C) data, we mapped identified eQTLs on genomic interactions reported in Mifsud et al. (Mifsud, et al. 2015). Anchors of promoter-promoter and promoter-other CHi-C interactions were analyzed for enrichment with the eQTLs (Supplementary Figure 22). The analysis shows that CHi- C anchors containing promoters are enriched with proximal eQTLs and depleted of the distal ones. Similar effect can be observed for ChIA-PET RNAPII anchors intersected with promoters (Supplementary Figure 22). However, unlike CHi-C anchors containing enhancers, ChIA-PET anchors intersected with enhancers are significantly enriched with distal eQTLs. The results for ChIA-PET data highlight the role of distal enhancers in gene regulation and may suggest that the interactions identified in RNAPII ChIA-PET are more transcriptionally active than the ones reported from CHi-C.

To state more firmly the relationship between proximal and distal eQTLs and chromatin activity, we collected (see Methods) sequencing (ChIP-seq) data for three histone modifications (H3K27ac, H3K4me3, H3K4me1) and information on chromatin accessibility (DNase-seq) and analyzed it in interaction anchors intersected with the eQTLs. H3K4me3 is primarily associated with promoters, H3K4me1 with active enhancers and H3K27ac with active promoters and enhancers (Ernst, Kheradpour, et al. 2011). As expected, interaction anchors altered by proximal eQTLs are enriched with promoter signal, whereas those affected by distal eQTLs with enhancer signal (Fig. 4i). This confirms that proximal eQTLs disrupt promoter-enhancer communication at the site of the promoter and distal eQTLs at the site of the enhancer. The results are statistically significant, even though interaction anchors are enriched with chromatin marks in general (Fig. 4i) (Tang, et al. 2015). Furthermore, eQTLs emerge in densely connected genomic regions (Fig. 4g and 4i). This is also reflected by the fact that a single eQTL often intersects more than one RNAPII interaction anchor (Fig. 4k).

We repeated the eQTL analysis described above for housekeeping genes only (selected based on Eisenberg et al. (Eisenberg and Levanon 2013)) to see if we find eQTLs for them and where the potential eQTLs are localized (see Methods). We found 36 unique eQTLs for 33 different housekeeping genes. None of the eQTLs is located within CTCF anchor, but we observe significant enrichment of them in RNAPII anchors (Fig. 4e). Therefore, there are differences in expression rates of housekeeping genes among samples and they are mainly correlated with alternations of long-range chromatin contacts mediated by RNAPII.

On the other hand, we separately analyzed immune-related genes as genes specific for the lymphoblastoid cell lines (see Methods). 14 eQTLs were identified for these genes, out of which 4 intersect CTCF anchors. Three of these are anchors which contain enhancers and 1 contains a promoter region.

Two of the immune-related eQTLs (deletion chr22:39357694-39388574 and CNV chr22:39359355-39379392) cover the same CTCF anchor (Supplementary Figure 23a). Both are eQTLs for genes APOBEC3A, APOBEC3B and CTA-150C2.16 (Supplementary Figure 23b), but the deletion completely excises APOBEC3B gene. In the presented samples (Supplementary Figure 23a) both of the SVs were identified, meaning that locus chr22:39357694-39388574 is (haplotype-specifically) excised in those genomes, which is reflected in CTCF signal for these samples.

Another example of an eQTL altering a CTCF anchor and regulating an immune-related gene is deletion chr17:73107713-73108273 (Supplementary Figure 24a). It is located over 750 kb apart from the correlated gene TRIM47 (Supplementary Figure 24b).

Whether cell-type-specific genes are distinguished targets for SVs altering core chromatin architecture (as CTCF is believed to form the backbone network of genomic interactions) is an interesting question. However, more extensive testing has to be done to explore this hypothesis.

### Modeling topological domains of individual genomes

We modeled structures of CCDs for all 2,504 individuals sequenced in the 1000 Genome Project at the resolution of individual chromatin loops, creating a catalogue of genomic topological variation of human population.

We focus on topological domains as they are believed to be the structural units regulating gene transcription by spatially isolating groups of enhancers and genes (Dixon, Selvaraj, et al. 2012), (Lupiáñez, et al. 2015), (Rao, et al. 2014), (Tang, et al. 2015), (de Laat and Denis 2013). In our opinion, these 3D models constitute a supportive insight into SV effects; their inspection can improve our understanding of functional impact and disease association of SVs.

Deletion of a CTCF binding site insulating the promoter of TAL1 gene from regulatory elements adjacent to the CMPK1 promoter was shown by CRISPR/Cas9 experiments to cause activation of TAL1, an oncogenic driver of T-cell acute lymphoblastic leukemia (Hnisz, et al. 2016) (Fig. 5a). 3D structures of the TAL1 locus generated with our algorithm illustrate fusion of the TAL1 promoter with the enhancer regions inside the insulated neighborhood formed as a consequence of the deletion (Fig. 5b). Euclidean distances calculated from the models quantify the accessibility of transcription-enhancing elements for the TAL1 promoter. The Euclidean distance between the promoter and a strong enhancer in the CMPK1 promoter and the mean distance between the promoter and enhancers located in the same CCD decrease significantly after the deletion (Fig. 5c). The models show how the promoter and the enhancer are brought even closer together within the insulated neighborhood by RNA-mediated chromatin interactions (Fig. 5d). This demonstrates that the models properly capture the features of the chromatin organization and that their inspection can give insights into functional consequences of SVs.

**Fig. 5.**
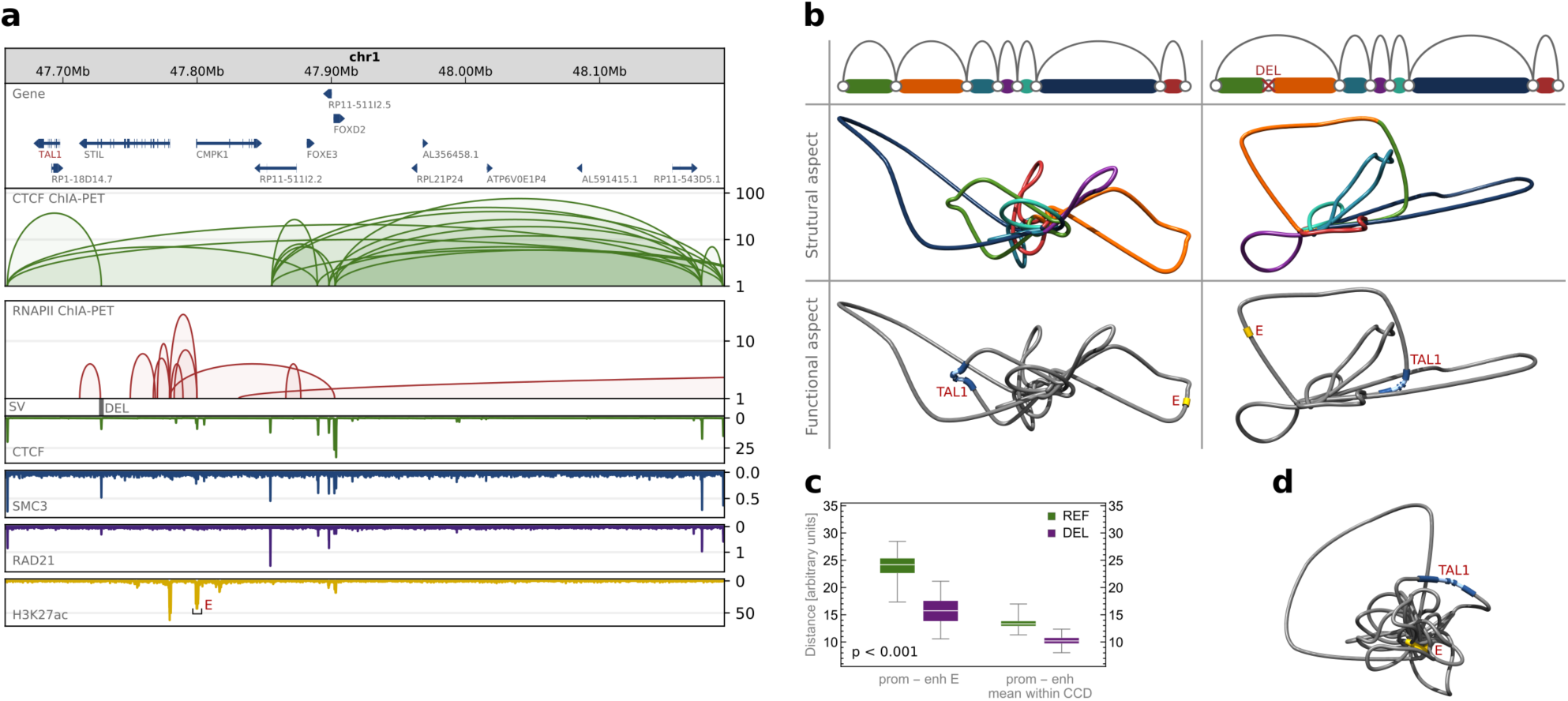
TAL1 locus. **a,** Browser view of a topological domain containing TAL1 gene and a deletion causing its activation. The deletion removes CTCF insulating the TAL1 promoter from enhancer (E). CTCF and RNAPII ChIA-PET interactions are shown along with ChIP-seq tracks for CTCF, cohesin subunits (RAD21 and SMC3) and H3K27ac which marks the enhancer E. **b,** Models presenting 3D structure of the TAL1 locus before and after introduction of the deletion removing CTCF. Chromatin loops forming the region have different colors, as presented on schematic drawings. Every picture has its replicate, in which TAL1 gene and enhancer E are marked. **c,** Euclidean distance between the TAL1 promoter and enhancer E and mean Euclidean distance between the promoter and enhancers located in the same CCD. In green, distribution of distances calculated in 3D models of the reference structure (REF), in magenta – in models with the deletion introduced (DEL). For each case 100 models were generated. The differences between REF and DEL groups are statistically significant (p-values much less than 0.001). **d,** 3D model of the TAL1 locus including RNAPII-mediated chromatin interactions.

As presented earlier, the inspection of a deletion in NFKBIA locus also led to functional interpretation of structural changes (Fig. 3c and 3e). More examples of expression-associated SVs disrupting long-range chromatin interactions can be found in the CHGV (Supplementary Figure 25) and inspected using the 3D-GNOME 2.0 – a web service providing the chromatin modeling method together with a visualization tool (http://3dgnome.cent.uw.edu.pl/).

## Discussion

We performed a genome-wide analysis of the impact of SVs on three-dimensional structure of the human genome. We showed that CTCF interaction anchors are depleted of SVs identified in genomes of healthy people. Moreover, anchors overlapped with enhancers or promoters occur to be conserved better than enhancers and promoters unrelated to CTCF interaction anchors. On the other hand, these genomic structural sites turned out to be enriched with SNPs previously associated with disease, and with autoimmune diseases in particular. This demonstrates the importance of these sites for proper genome functioning. Unexpectedly, CTCF binding sites involved in insulation of topological domains, occur not to be particularly conserved, and are even slightly enriched with duplications. Duplications of CTCF anchors at domain boundaries could be a protective and evolutionary mechanism. On the other hand, the fact that they are often targeted by other SVs suggests that they can also be important elements used in genome regulation.

Even though interacting genomic segments seem to be evolutionarily protected from mutation, we still observe interesting examples of SVs disrupting long-range chromatin contacts and correlated with transcription rates of genes located within the same topological domain. Even a single point mutation can be sufficient to make an interaction anchor active or inactive and alter chromatin looping and gene expression as a consequence. This shows the explanatory potential of this type of 3D genome studies.

Examples of expression-related SVs in CTCF interaction anchors are not abundant. Genome-wide analysis demonstrates that eQTLs are rarely located in CTCF anchors or CCD borders. Instead they favorably alter RNAPII interaction anchors, which strongly suggests that structural variation influences transcription of genes mainly by directly altering long-range transcription-related chromatin contacts. We observe cases in which both CTCF and RNAPII anchors are modified in a CCD to regulate gene transcription, but the overall analysis indicates that the chromatin structure built by CTCF is strongly conserved across individuals and variation in gene transcription occurs by modifications of RNAPII interactions formed within this structure. This hypothesis is consistent with the model presented by Tang et al. (Tang, et al. 2015). However, we suspect that the fact that we did not identify many examples of eQTLs located in CTCF anchors may also mean that the linear model used to detect eQTLs is too simple to account for complex nonlinear changes in gene transcription caused by modification of CTCF-mediated chromatin looping. This requires further investigation.

Together with analyzing the impact of SVs on genomic structural elements we devised a method for modeling changes that SVs introduce in 3D chromatin structure. We also prepared a first catalogue of 3D structures of CCDs for all 2,504 individuals sequenced in the 1000 Genome Project at the resolution of individual chromatin loops, which represent the topological variability of the human genome. We believe that further work on computational prediction of the chromatin 3D structures based on different factors changing among individuals is required, as the emerging evidence shows that chromatin structure is a crucial element to understand the genome regulation.

## Methods

### Genomic interactions

Genomic interactions analyzed in this study are 80,157 CTCF PET clusters and 100,263 RNAPII PET clusters identified by Tang et al. (Tang, et al. 2015) for the GM12878 cell line. We refer the reader to this work for details on data processing pipeline used to find these interactions. Briefly, pair-end reads (PETs) sequenced in long-read ChIA-PET experiment were mapped to the human reference genome (hg19). Inter-ligation PETs were selected by the criterion of genomic span between the two ends of a PET exceeding 8 kb. Inter-ligation PETs overlapping at both ends were clustered together creating unique contacts (PET clusters) between 2 specific interaction loci, of strength equal to the size of the cluster. PET clusters of strength smaller than 4 were discarded.

### ChIP-seq consensus peaks

We analyzed the directionality of CTCF interactions similarly to Tang et al. (Tang, et al. 2015). CTCF, RAD21 and SMC3 uniform ChIP-seq peaks available for the GM12878 cell line were downloaded from the ENCODE database (The ENCODE Project Consortium 2012). We extracted the 4-way consensus regions from all 4 sets of CTCF peaks to get highly credible CTCF-binding peaks. The same consensus was performed on RAD21 and SMC3 ChIP-seq segments to identify cohesin-binding peaks. Finally, 24,118 consensus regions from the CTCF and cohesin consensus peaks were obtained.

### CTCF motif identification

We searched the CTCF-cohesin consensus peaks for CTCF-binding motifs. Nucleotide sequence of each of the CTCF-cohesin peaks was extracted from the hg19 assembly deposited at http://genome.ucsc.edu using BEDTools getfasta utility (Quinlan and Hall 2010) and provided as an input to STORM (Schones, Smith and Zhang 2007). Given the position weight matrix of a particular transcription factor-binding motif, STORM predicts the occurrences of the factor-binding motifs in provided DNA sequences. We performed the search with CTCF position weight matrix MA0139.1 downloaded from the JASPAR database (Khan, et al. 2018) and found CTCF-binding motifs in 22,677 out of the 24,118 CTCF-cohesin consensus peaks. Only motifs having score higher than 0 were considered as valid and for each peak a motif with the highest score was selected.

### Assigning orientation to CTCF loops

The CTCF motifs were overlapped with CTCF PET clusters. 42,177 out of the 80,157 CTCF PET clusters had both anchors overlapped by CTCF-cohesin consensus peaks and only 2,806 (4%) of them had no intersections with the consensus peaks at neither of sides. For 38,196 out of the 42,177 clusters (91%) at least one CTCF motif was found at both anchors. 31,422 of these had exactly one motif at either side. PET clusters with anchors having more than one CTCF motif and of contradictory orientations were filtered out. From the 35,180 CTCF PET clusters with motifs of unique orientation in both anchors, 16,239 (46%) had motifs of convergent orientation at the two anchors, 12,195 (35%) had motifs of tandem right orientation, 4,090 (12%) PET clusters were of tandem left orientation and 2,656 (7%) were of divergent orientation.

### Chromatin contact domains

In this study, we used 2,267 CTCF-mediated chromatin contact domains (CCDs) identified by Tang et al. (Tang, et al. 2015). We refer the reader to this work for the details of CCD calling. Briefly, CCDs were identified by searching each chromosome for genomic segments continuously covered with CTCF PET clusters supported by CTCF-cohesin consensus peaks. Each identified CCD starts where the most upstream CTCF anchor from all the anchors of the CTCF PET clusters comprising the CCD starts, and ends where the most downstream CTCF anchor ends. To define borders of the CCDs more accurately, CTCF motifs found in the CTCF-cohesin consensus peaks and positioned within outermost anchors were identified. From these, outermost CTCF motifs were selected as CCD borders. Genomic regions complementary to CCDs (less hg19 reference genome assembly gaps) were defined as CCD gaps.

### Enhancers and promoters

Definitions of enhancers used throughout this study were extracted from ChromHMM (Ernst and Kellis, ChromHMM: automating chromatin state discovery and characterization 2012) hg19 annotations for the GM12878 cell line downloaded from the ENCODE database (The ENCODE Project Consortium 2012). Both weak and strong enhancer annotations were adopted. Promoters were defined as ±2 kb regions surrounding gene transcription start sites (TSSs). The TSS coordinates were adopted from the GENCODE release 27 (mapped to hg19) (Harrow, et al. 2012). Only promoters for protein-coding genes were considered. Promoters were defined as active if overlapped with anchors of RNAPII PET clusters or with RNAPII consensus ChIP-seq peaks and defined as inactive otherwise. RNAPII consensus peaks were obtained by performing consensus on 3 sets of RNAPII uniform ChIP-seq peaks available for the GM12878 cell line in the ENCODE database.

### Enrichment analyses of SVs in genomic elements

In this study, we tested various genomic elements for enrichment or depletion with structural variants (SVs). These tests were conducted according to a common scenario. Genomic elements of a given type represented by their positions in the hg19 reference genome were intersected with the positions of SVs. The ones having at least 1 bp overlap with at least 1 SV were counted. The genomic elements were then intersected with simulated SVs from 1,000 sets generated by randomly shuffling positions of the original SVs, and the null distribution of counts of genomic elements overlapped with SVs was calculated. Each shuffled set contained the same number of elements in total and the same number of elements in subsets (deletions, duplications, etc.) as the real SV set. Elements of these sets were equally distributed on chromosomes and equally distributed in length to the real SVs. The operations of shuffling and intersecting genomic segments were performed with BEDTools (Quinlan and Hall 2010). The enrichment or depletion of genomic elements overlapped with the real SVs compared to the same elements overlapped with randomly positioned simulated SVs was expressed as log2 fold change of the number of the former versus the mean of the distribution of the number of the latter. Values of the measure were represented by the height of bars in the plots. Error bars in the plots show the standard deviations of log2 fold changes. To estimate the statistical significance of the test results, p-values were calculated and marked above the bars by stars (3 stars – p-value < 0.001, 2 stars – p-value < 0.01, 1 star – p-value < 0.1).

### Subsets of SVs in enrichment analyses

For individual tests the set of SVs was divided into various subsets. In particular, SVs with variant allele frequency (VAF) lower than 0.001 were considered separately in certain tests. In others, SVs were grouped according to the ancestry of individuals they were identified in and sets of SVs emerging uniquely in one subpopulation were also created. The special set of SVs correlated with gene expression (eQTLs) was subdivided into 2 sets: a set of eQTLs located closer on the DNA chain than 17,800 bp apart from genes they modified, and a set of eQTLs located further apart from their genes. The distances were calculated between TSSs (as defined in GENCODE version 12) and centers of eQTL segments.

### Subsets of GWAS SNPs

The set of GWAS SNPs used in this study was derived from the NHGRI-EBI GWAS Catalog, version from January 31, 2018 (MacArthur, et al. 2017). SNPs of traits associated with autoimmune diseases and hematological parameters were extracted as separate sets and mapped to dbSNP Build 150 for hg19 human genome assembly. SNPs mapping outside the main chromosome contigs, not having dbSNP ID or without coordinates on the hg19 and records containing multiple SNPs were excluded. This resulted in 2,330 and 3,919 unique SNPs associated with autoimmune diseases and hematological parameters respectively. For permutation tests with SNPs identified in healthy samples in the 1000 Genomes Project, we extracted a random sample of 1 million elements from the whole set of SNPs to limit the computation time and storage space.

### Genomic elements in enrichment analyses

Analyzed in permutation tests genomic elements associated with genes (annotated protein-coding sequence regions (CDSs), untranslated regions (UTRs) in protein-coding regions, exons and introns) were adopted from the GENCODE release 27 (mapped to hg19). Permutation tests with eQTLs were an exception – in this case, gene elements from version 12 of GENCODE were used to maintain the consistency with the expression data which was analyzed with the earlier version of GENCODE. The positions of transcription factor binding sites (TFBSs) were adopted from a file with uniform TFBS peaks downloaded from ENCODE (http://genome.ucsc.edu/cgi-bin/hgTrackUi?db=hg19&g=wgEncodeRegTfbsClusteredV3). In permutation tests, CTCF anchors supported and not supported by cohesin peaks were also analyzed. Anchors supported by cohesin peaks are anchors of CTCF PET clusters, which showed at least 1 bp overlap with cohesin-binding consensus peaks (identified as described above).

### GM12878 haplotype-specific ChIA-PET interactions

Phased ChIA-PET interactions for GM12878 cell were taken from the work of Tang et al. (Tang, et al. 2015) and we refer the reader to this article for details of the ChIA-PET data phasing analysis. Briefly, a ChIA-PET read was assigned to the maternal or paternal haplotype if a nucleotide in its sequence aligned with a GM12878 heterozygous SNP matched the respective allele. All the reads from the two ends of all inter-ligation PETs were distributed between the haplotypes in this manner, yielding PETs of 6 possible types: M-M, P-P, M-N, P-N, M-P and N-N, where M stands for maternal, P for paternal and N for not determined haplotype of an anchor. Following the outcomes of the analyses performed by Tang et al. we treat both M-M and M-N PETs as maternal interactions and both P-P and P-N PETs as paternal interactions. There are 917 maternal and 1233 paternal interactions. The phased SNPs used for the analysis were downloaded from ftp://gsapubftp-anonymous@ftp.broadinstitute.org/bundle/2.8/hg19/.

### Analysis of CTCF interaction anchors altered by SNPs

To test impact of SNPs on the probability of CTCF binding to a CTCF anchor, we searched the nucleotide sequence of the anchor for CTCF motifs and compared the number and scores of these motifs with the CTCF motifs identified in the nucleotide sequence of this anchor after introduction of alternative alleles. Only motifs with score higher than 0 were taken into consideration. Identification of CTCF motifs was performed as described in the section *CTCF motif identification* above.

We used ggseqlogo R package (Wagih 2017) to generate sequence logos from the frequency matrix MA0139.1 downloaded from the JASPAR database.

### mRNA quantifications

PEER normalized expression levels of 23,722 genes provided by gEUVADIS consortium were used in our analyses. We refer the reader to Lappalainen et al. (Lappalainen, et al. 2013) for details on the process of transcriptome quantifications. In short, RNA-seq read counts over genes annotated in GENCODE (version 12) were calculated. This was done by summing all transcript RPKMs per gene. Read counts were corrected for variation in sequencing depth by normalizing to the median number of well-mapped reads among the samples and for technical noise. The latter was removed using PEER (Stegle, et al. 2012). We logarithmized the corrected quantifications and standardized the distributions of transcription rates (for each gene individually).

### Genotypes

Definitions of genomic sequence variations were taken from Sudmant et al. (Sudmant, et al. 2015). This SV set is a refined version of the callset released with the 1000 Genomes Project marker paper (The 1000 Genomes Project Consortium 2015). Only SVs (deletions, duplications, copy number variants, inversions and insertions) were considered, SNPs were not included in the analysis. The genotype of an individual was represented as a sum of SV copies present on homologous chromosomes of the individual. Deletions were indicated by negative numbers. For example, if an individual had a deletion on both copies of a chromosome, the genotype was −2. If it had 2 more copies of a genomic region (in relation to the reference genome) on one chromosome from the pair and 1 additional copy of this region on the second chromosome from the pair, the genotype was 3. Genotypes unchanged compared to the reference sequence (hg19 in this case) had codes 0. Genotypes of abundance lower than 1% in the studied population were neglected.

### Linear models

gEUVADIS consortium provides RNA-seq data for 462 samples, but only a subset of these (445 samples) was genotyped in the 1000 Genomes Project. Thus, our analyses were performed on a population of 445 individuals for which both transcription and genotype data was available. Only genotypes of abundance higher than 1% were considered. Sex chromosomes were excluded from the analyses. We took the logarithms of the PEER-normalized expression levels for calculations to correct it for far outliers and standardized the data. We started the analysis by performing Principal Component Analysis (PCA) in the 23,722-dimensional space of gene expression rates. Based on the Scree Plot (Supplementary Figure 26) we decided to keep first 100 principal components. Only genes having contributions to these components not smaller than 0.01 were considered in the further analyses. By running this procedure, we got 14,853 genes of largest contribution to the variance in gene transcription between samples. Every SV lying in the same CCD as one of these genes was tested for being eQTL for this gene. Least-squares linear regression between expression rates and genotypes was performed for each gene-SV pair. The slopes of the linear models were tested for statistical significance. First, for each linear model two-sided p-value was calculated in test with null hypothesis that slope is 0 (Wald Test with t-distribution of the test statistics). Second, for each gene we permutated the expression rates relative to genotypes 1,000 times, recalculating at each iteration the linear regression for each gene-SV pair and recording the minimal p-value among all pairs. Adjusted p-values were calculated for each gene by dividing ranks of the observed p-values in the list of p-values obtained in permutations, by the number of permutations. Finally, to correct for multiple testing across genes, we applied the Benjamini-Hochberg procedure to the adjusted p-values, estimating q-values. At FDR 0.1 we found 203 genes with eQTLs. The same procedure was employed to identify eQTLs for housekeeping genes, except that PCA step was omitted. By mapping the names of housekeeping genes reported in Eisenberg et al. (Eisenberg and Levanon 2013) on GENCODE (version 12), we obtained a list of 3,784 genes. We found eQTLs for 27 of them. A complete list of all discovered eQTLs is provided in Supplementary Table 2.

### Immune-related genes

Names and coordinates on the hg19 assembly of immunity genes were downloaded from InnateDB (Breuer, et al. 2013). The gene names were mapped on GENCODE (version 12) for the purpose of the eQTL analysis. The final gene set contained 1051 elements.

### ChIP-seq signal tracks

Raw sequencing data from CTCF ChIP-seq experiments published by Kasowski et al. (Kasowski, et al. 2013) was processed to obtain CTCF signal tracks for 10 lymphoblastoid cell lines (GM12878, GM10847, GM12890, GM18486, GM18505, GM18526, GM18951, GM19099, GM19238, GM19239). The sequencing reads were aligned to the hg19 assembly using Bowtie2 aligning tool (Langmead and Salzberg 2012). The alignments were then passed to the bamCoverage utility from the deepTools2.0 toolkit (Ramírez, et al. 2016) to obtain RPM values genome-wide (the following command was evoked: bamCoverage -b input.bam -o output.bw -of bigwig - -binSize 10 --numberOfProcessors max/2 --normalizeUsing CPM -- ignoreForNormalization chrX --extendReads --samFlagInclude 64). Sequencing reads for every sample and for every experimental replicate were processed separately. Signals prepared for different biological replicates but for the same sample, were merged to an averaged signal using the mean operator from the WiggleTools1.2 package (Zerbino, Johnson, et al. 2014). Each CTCF signal track included in the figures presents RPM values for a particular genomic region divided by the maximal value of the signal in this region.

Presented RNAPII signals were downloaded from the UCSC database (https://genome.ucsc.edu/cgi-bin/hgFileUi?db=hg19&g=wgEncodeSydhTfbs). The signal values were also max-normalized for the comparison purposes.

### Phased ChIP-seq signal tracks

Haplotype-specific CTCF ChIP-seq signals for 10 lymphoblastoid cell lines (GM12878, GM10847, GM12890, GM18486, GM18505, GM18526, GM18951, GM19099, GM19238, GM19239) were obtained from the raw sequencing data published by Kasowski et al. (Kasowski, et al. 2013). The reads sequenced for a particular cell line were aligned with Bowtie2 aligning tool to the individualized nucleotide sequences of maternal and paternal chromosomes of this cell line. Only perfectly aligned reads were considered as valid. The sequences of maternal and paternal chromosomes were prepared with the vcf2diploid tool from the AlleleSeq pipeline (Rozowsky, et al. 2011) using SNP phasing information from the phase 3 of the 1000 Genomes Project. Separate ChIP-seq signals for maternal and paternal chromosomes of the individual samples were calculated from the alignments prepared for the respective chromosomes analogously to the non-phased signals.

### Modeling three-dimensional chromatin structures with 3D-GNOME

The ChIA-PET datasets typically consist of two types of interactions: high-frequency PET clusters (in the order of tens of thousands) representing strong, specific chromatin interactions and singletons, numerous (in the order of tens of millions), but representing mostly non-specific and spurious contacts.

To make the best use of the information carried by these two distinct types of contacts we employed a multiscale approach: first we used the singletons to guide the low-resolution, megabase-scaled modeling, and then we used PET clusters to refine the obtained structures, achieving resolutions up to a few kilobases. We note that this approach is consistent with the widely accepted model of genome organization, in which main roles are played by topological domains and chromatin loops. Here, at the stage of the low-resolution modeling we attempt to position the topological domains relative to each other, and in the high-resolution we model the position and shape of individual chromatin loops.

### Low-resolution (chromosome) level modeling

The structure of a chromosome is represented using a ‘beads on a string’ model. First, the chromosome is split into a number of approximately megabase-sized regions. Ideally, each region would correspond to a single topological domain. In practice, the split is made based on the patterns of PET clusters interactions (as a consequence, different regions typically will have different lengths. See (Szałaj, et al. 2016) for details of the procedure). Next, singleton heatmaps are created (much like the widely used Hi-C heatmaps, but with unequal bins). We treat an interaction frequency *f*_*ij*_ between a pair of regions *i* and *j* as a proxy of 3D distance *d* _*ij*_ between corresponding beads, assuming an inverse relationship 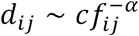, with *α* being a scaling exponent, and use Monte Carlo simulated annealing to position the beads to minimize the energy function 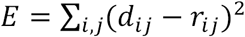, where *r*_*ij*_is the actual distance between beads corresponding to regions *i* and *j.*

### High-resolution modeling

In the second step we model the shape and position of chromatin loops inside a single domain. We begin by splitting the interaction network given by PET clusters contained within a domain into a number of disjoint connected components that we call *blocks*. This allows us to model blocks independently. The modeling of each block is carried out in 2 steps. First, in the anchor step, we position the anchors of the loops identified by ChIA-PET relative to each other. The preferred distance between a pair of anchors *i* and *j* connected by a loop depends on the frequency of PET cluster solely and is given by 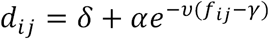, where *δ, α, ν* and *γ* are all parameters (if the anchors are not connected, then *d*_*ij*_ is not specified). The energy function is identical in the form to the one used at the chromosome level, and we again use Monte Carlo simulations to find the optimal arrangements of the anchors. Then, in the subloop step, we keep the anchors positions fixed, and we try to model loops so that their shape, as well as relative position to other loops best fit both the data and the physical constraints. Each loop is represented by *k* subanchor beads inserted between neighboring anchors. We define stretching and bending energy terms as 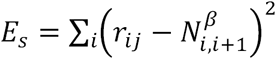 and 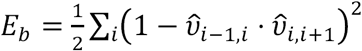, where *N*_*j,j*+1_ is a genomic distance between anchors *j* and *j+1*, 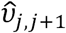 is a unit vector pointing from anchor *j* to *j+1*, and *β* is a constant parameter. To model the influence of short-range singleton interactions we calculate the expected distances between all subanchor beads given only the physical constraints, and then modify these distances based on the high-resolution singleton heatmaps for blocks. These updated distances are used in the third energy term *E*_*h*_ *=* ∑ _*i,j*_*(d*_*ij*_ - *r*_*ij*_)^2^, with *d*_*ij*_ and *r*_*ij*_ defined as previously, but now for subanchor beads. The total energy function is simply *E*_*s*_ + *E*_*A*_ + *E*_*F*_, and again, the Monte Carlo simulation are used for the optimization.

### Modeling impact of SVs onto the three-dimensional chromatin structure

The algorithm modifies the reference genomic interactions and topological domains introducing information on SVs. The resulting genomic interaction data is then passed to the 3D-GNOME modeling engine to obtain predicted 3D structures adjusted for SVs.

Anchors intersected by deletions are removed from the interaction pattern of a topological domain being modeled. As a result, all the interactions stemming from these anchors are eliminated yielding a structure with fewer loops, and with the loops directly neighboring the deletion being shorter or longer depending on the interaction pattern and the deletion size. The anchors intersected by duplications are duplicated along with all the contacts they have with other genomic segments. This rule reflects the potential of the newly introduced anchor to interact with the rest of the chromatin chain analogously to the duplicated site. Introducing duplications also results in elongation of loops overlapping the duplicated site. Inversions of genomic segments containing anchor sites result in changing positions of the anchors and its directionality relative to other anchors. The algorithm attempts to arrange orientations of interacting CTCF motifs concordantly, which reflects the preference of CTCFs to have symmetric conformation in dimers (Tang, et al. 2015). The insertions detected in the 1000 Genomes Project are almost solely insertions of transposable elements, which do not introduce new CTCF binding sites to the genome. Nevertheless, our algorithm enables introducing new CTCF binding sites to domain structures. Along with such insertions, new contacts are introduced between the inserted anchors and their neighbors.

The algorithm accounts also for SVs that miss the CTCF binding sites, but introduction of these results only in shortening or extending the corresponding chromatin loops.

### Comparison of CTCF interaction segments among different individuals

In order to assess fraction of CTCF interaction segments conserved among different lymphoblastoid cell lines, we calculated a number of CTCF anchors identified in GM12878, which were intersected with CTCF ChIP-seq peaks called in the remaining lymphoblastoid cell lines.

Since the available sets of CTCF ChIP-seq peaks for multiple lymphoblastoid cell lines (Kasowski, et al. 2013) differed highly in size (Supplementary Figure 27), we performed an additional filtering on them. Only those CTCF ChIP-seq peaks which intersected with consensus CTCF binding sites were selected for each of the lymphoblastoid cell lines. The consensus CTCF binding sites were collected from the Ensembl 92 database (Zerbino, Achuthan, et al. 2018). They were identified by: first, performing a genome segmentation based on a variety of genome-wide assays from multiple cell types (including histone modification ChIP-seqs, TF ChIP-seqs, DNase-seq) and selecting segmentation state corresponding to CTCF peaks; and second, annotating the position of CTCF binding sites within the peaks using JASPAR position weight matrix MA0139.1. For more details, see Ensembl website.

The consensus CTCF binding sites were downloaded via BioMart interface (www.ensembl.org/biomart) in genomic coordinates of hg38 assembly and converted to hg19 coordinates using UCSC liftOver tool.

Comparable datasets were obtained by the filtering (Supplementary Figure 28).

### Aggregate analysis of ChIP-seq signals in altered interaction anchors

In order to analyze the overall behavior of ChIP-seq signal in interacting genomic segments affected by deletions or duplications, we performed an aggregate analysis. The same procedure was applied regardless of SV type (deletion or duplication) and ChIP-seq target protein (CTCF or RNAPII). We describe it using an example of CTCF interacting segments affected by deletions.

First, CTCF anchors intersected by deletions exhibited by at least one of the 10 lymphoblastoid cell lines with available CTCF ChIP-seq data (Kasowski, et al. 2013) were identified. For each such anchor 200 bins were defined – 100 bins of equal size covering an anchor and 50 equal-size bins covering genomic regions 500 bp upstream and downstream from the anchor. Averaged raw CTCF ChIP-seq signal was calculated in every bin for every sample. For each sample a mean of the signal over the whole genome was found and subtracted from the extracted binned signal values. The maximal mean from all the samples was then added to the values to make them positive. Obtained values were then averaged over the samples exhibiting and not exhibiting the deletion. The log2 of the ratio of the signal values obtained for the first group to the ones obtained for the second group was calculated. The log2 fold changes calculated for all the anchors affected by deletions were then averaged and plotted in Supplementary Figure 7a.

### Histone marks and DNase-seq data

H3K27ac, H3K4me3, H3K4me1 and DNase-seq data for GM12878 was downloaded from the ENCODE database in a form of bigWig files containing signal fold change over control. The data on histone marks was generated by the laboratory of Bradley Bernstein. The DNase-seq was performed by Gregory Crawford’s group.

## Acknowledgements

This work was carried out within the TEAM program of the Foundation for Polish Science co-financed by the European Union under the European Regional Development Found, co-supported by Polish National Science Centre (2014/15/B/ST6/05082) and the grant 1U54DK107967-01 “Nucleome Positioning System for Spatiotemporal Genome Organization and Regulation” within 4D Nucleome NIH program.

## Author Contributions

D.P. and M.S. conceived the methodology of modeling CCDs of individual genomes at the population scale. M.S. implemented the algorithm. P.S., D.P., Z.T., Y.R. devised the 3D-GNOME used as a main engine for modeling. M.W. extended the 3D-GNOME web service to provide the SV-including modeling method (3D-GNOME 2.0). M.S. designed the statistical analysis part. M.S. and A.K. performed the analyses. M.S., A.K., Z.T. extracted and prepared data for the analyses. M.S., P.S., A.K., Z.T. and Y.R., D.P. prepared the manuscript. D.P., Y.R., Z.T. supervised the study.

## Additional Information

3D-GNOME 2.0 - a web service implementing the chromatin modeling method with SV integration is available at: http://3dgnome.cent.uw.edu.pl/.

